# Multiple-kernel learning for genomic data mining and prediction

**DOI:** 10.1101/415950

**Authors:** Christopher M. Wilson, Kaiqiao Li, Pei-Fen Kuan, Xuefeng Wang

## Abstract

Advances in medical technology have allowed for customized prognosis, diagnosis, and personalized treatment regimens that utilize multiple heterogeneous data sources. Multiple kernel learning (MKL) is well suited for integration of multiple high throughput data sources, however, there are currently no implementations of MKL in R. In this paper, we give some background material for support vector machine (SVM) and introduce an R package, RMKL, which provides R and C++ code to implement several MKL algorithms for classification and regression problems. The provided implementations of MKL are compared using benchmark data and TCGA ovarian cancer. We demonstrate that combining multiple data sources can lead to a better classification scheme than simply using a single data source.

## 1 Introduction

Integrating multiple heterogeneous high throughput data sources is an emerging topic of interest in cancer research. Making decisions based upon metabolomic, genomic, etc. data sources can lead to better prognosis or diagnosis than simply using clinical data alone. Support vector machines (SVM) are not suitable for analyzing multiple data sources in a single analysis. SVM employs the kernel trick, thus it is able to construct nonlinear classification boundaries. However, obtaining an optimal kernel type, hyperparameter, and regularization parameter are challenging tasks. Cross validation can be employed to make these selections. Ultimately, there may not be one single optimal kernel, but rather a convex combination of several kernel representations of the data. Methods that produce a classification rule based on a convex combination of candidate kernels are referred to as multiple kernel learning (MKL) methods.

We present an R package, RMKL, which can implement cross validation for training SVM and support vector regression models, as well as MKL for both classification and regression problems. Three implementations of MKL classification are included, SimpleMKL proposed by Rakotomamonjy *et al.* (2008), Simple and Efficient MKL (SEMKL) proposed by Xu *et al.* (2010), and SpicyMKL (DALMKL) presented by Suzuki *et al.* (2011). Each of these implementations were presented in MATLAB, but to our knowledge RMKL is the first package that implements MKL in R.

We provide a brief summary of the mathematical theory of SVM and MKL in section 2. In section 3, we describe the features included in our package RMKL. In section 4, we illustrate RMKL using benchmark datasets and predict survival outcomes using TCGA Ovarian dataset with clinical and miRNA data. Finally, in section 5, we make a few closing remarks.

## 2 Background

### 2.1 Support Vector Machine

We will be considering samples (*x*_*i*_, *y*_*i*_), where the outcome *y* = {−1, 1} and the vector of covariates *x* ∈ *χ*. The goal of SVM is to find the hyperplane, {*w* : *w · x* + *b*,} that correctly separates the data into two classes and has the largest possible largest distance between the boundary of the two groups, which is referred to as margin. The SVM classification rule is defined to be *y*(*x*) = *w · x* + *b*. The SVM problem can be expressed as the following convex optimization problem:

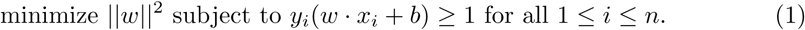

Note that *y*_*i*_(*w · x*_*i*_ + *b*) greater than 1 if *y*_*i*_ and *w · x*_*i*_ + *b* have the same sign, i.e. the sample is correctly classified. This problem is known as the hard margin formulation and is only feasible when two groups can be perfectly separated by linear function.

It is rare that data can perfectly linearly separable. We can relax (1) so that samples are allowed to be misclassified, by incorporating a penalty for samples that misclassified. The following optimization convex problem is referred to as the soft margin problem:

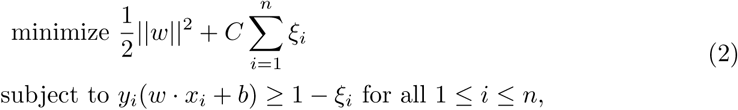

where *ξ*_*i*_ = max(0, *y*_*i*_(*w · x*_*i*_ + *b*)) and is known as the hinge loss function. The parameter *C* controls the penalty of misclassification, and a value for *C* is typically found via cross validation. Larger values of *C* can lead to a smaller margin to minimize the misclassifications, while smaller values of *C* may produce a larger margin that can lead to more misclassifications. Problem (2) is typically not solved directly, but rather by solving the Lagrangian dual.

The Lagrangian is the sum of the original objective function and a term that involves the constraints and multiplier. The Lagrangian of (2) is given below:

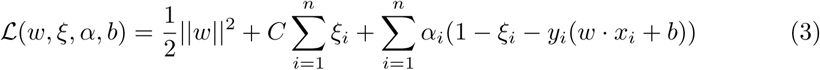

where *α*_*i*_ ≥ 0. The minimizers of *ℒ* are found by setting the gradient of *ℒ* equal to zero and solving the resulting system of equations:

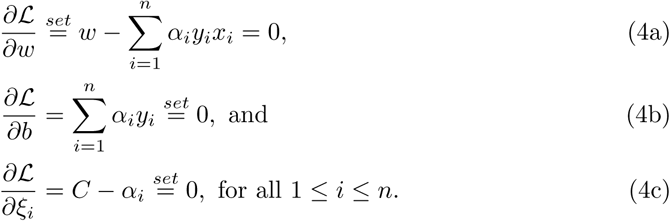

Equations (4b) and (4c) provide two new constraints, namely 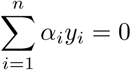 and *α*_*i*_ ≤ *C*, and provides a representation for the optimal hyperplane 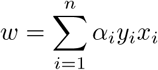 Plugging the solutions of (4) into the Lagrangian (3) yields the dual problem:

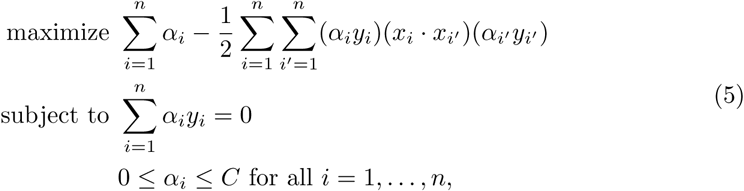

where (*x*_*i*_ *x*_*i′*_) denotes the dot product between *x*_*i*_ and *x*_*i′*_ This problem is a quadratic programming problem that can solved with many solvers and can be solved much more efficiently than (2). The Karush-Kuhn-Tucker (KKT) conditions are necessary conditions for solving a non-linear programming problem and for SVM these conditions are:

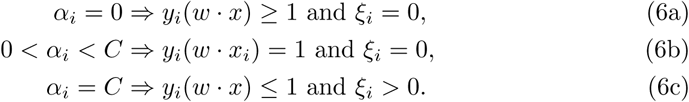

The resulting classification function produced by SVM algorithms is computed with only consider samples where 0 < *α*_*i*_ < *C*, which correspond to points that are on the margin, second condition. These points are called support vectors and the number of support vectors is can be much smaller than the number of samples which helps make SVM algorithms faster.

If data are not linear separable, kernels can be employed to map data into a higher dimensional feature space where the data are linearly separated. A kernel function *K*: *𝒳 × 𝒳 →* ℝ that for all *x*_*i*_, *x*_*i′*_ that satisfies *K*(*x*_*i*_, *x*_*i′*_) := (*ϕ* (*x*_*i*_) · *ϕ* (*x*_*i*_*I)*) where *ϕ*: *χ* → *ℋ*,and *ℋ* and is a Hilbert space. Kernel functions are different similarity measures between samples and *K* is a symmetric positive definite matrix which aid in solving optimization problems that we will introduce. The above derivation can be extended to non-linear classification by simply replacing *w* and *x*_*i*_, in (2) and (3), *f* (*x*) with *ϕ* (*x*_*i*_) and *f* (*x*) = *K*(*·, x*) respectively, where which yields

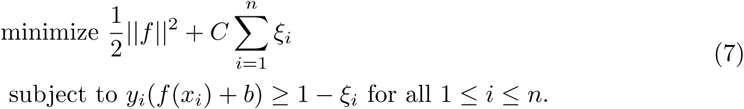

The Lagrangian dual can be constructed in a similar fashion as above, of (7) which yields:

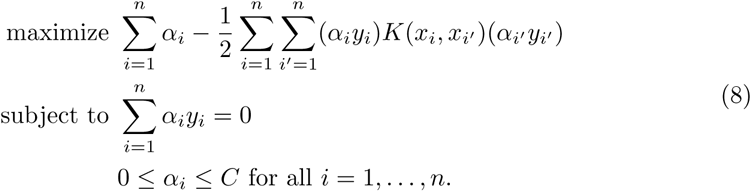

Since **K** is a symmetric positive definite matrix, (8) is a quadratic programming problem. Unlike SVM, we there is typically not a closed form expression for *f* (*x*), thus it is difficult to interpret the results.

Kernel selection is crucial and typically impossible, to confirm visually if input space has more than two dimension. Selecting an inappropriate kernel may lead to overfitting, or lead to a classification rule that misses important characteristics in the input space. Common kernels that are used for continuous predictors include:

1.Linear Kernel: *K*(*x*_*i*_, *x*_*i′*_) = (*x*_*i*_ · *x*_*i′*_)

2.Polynomial Kernel: *K*(*x*_*i*_, *x*_*i′*_) = (*γ \** (*x*_*i*_ *· x*_*i′*_) + *ν*)^*d*^ where *γ, ν* ∈ ℝ, *d* ∈ ℤ^+^

3.Gaussian Kernel: *K*(*x*_*i*_, *x*_*i′*_) = exp{*-σ ‖ x*_*i*_ *- x*_*i′*_ *‖*^2^}, where *σ >* 0

4.Sigmoid Kernel: *K*(*x*_*i*_, *x*_*i′*_) = tanh(*γ*(*x*_*i*_ · *x*_*i*_*′)* + *ν*), where *γ, ν* ∈ ℝ

Gaussian, polynomial and sigmoid kernels have parameters embedded within them, *σ* controls the radius of the Gaussian kernel. A scale, *γ*, and constant term, *ν*, can be specified for polynomial and sigmoid kernels. The selection of internal parameters is also vital to the success of the classification rule. For instance, the Gaussian kernel with a large radius can be similar linear or polynomial classification rule. A disadvantage of a Gaussian kernel is that can it lead to overfitting compared to simpler kernels such as linear or polynomial kernels.

Typically, medical studies include clinical variables, such as patient demographic characteristics as predictors and the aforementioned kernels are not appropriate for categorical predictors. Daeman, suggested the following kernel:

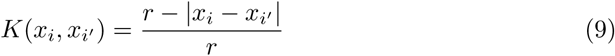

for each ordinal or continuous demographic factors, and

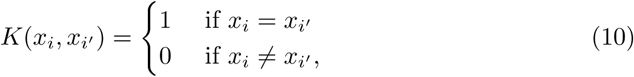

for nominal demographic factor. These kernels should be applied to each individual demographic variable and combined using a global measure of similarity of two samples by averaging each of the kernel values.

### 2.2 Multiple Kernel Learning

It has been shown that the convex combinations of kernel functions is a kernel function. An avenue for improvement is to utilize several different representations of the data and allow an algorithm use a weighted average of these representations of the data. This can help automate kernel selection by using a combination a kernel functions for a set of candidate kernels, this is the main idea of multiple kernel learning (MKL). Combining kernels is possible by decomposing the input space into blocks as follows *χ*=*χ*_1_ ×· · ·× *χ*_*m*_ where each sample can be expressed as *ϕ* (*x*_*i*_) = (*ϕ* _*i*_(*x*_*i*1_),*…, ϕ* _*i*_(*x*_*im*_)).MKL can be formulated as the following optimization problem:

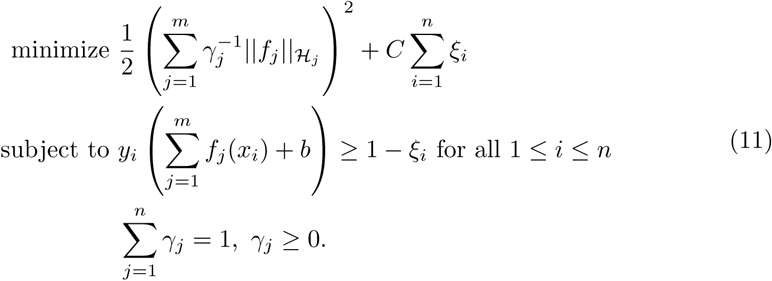

This problem remains convex, however it is not smooth which leads to computational issues. Using the same procedure as before, the Lagrangian dual given below:

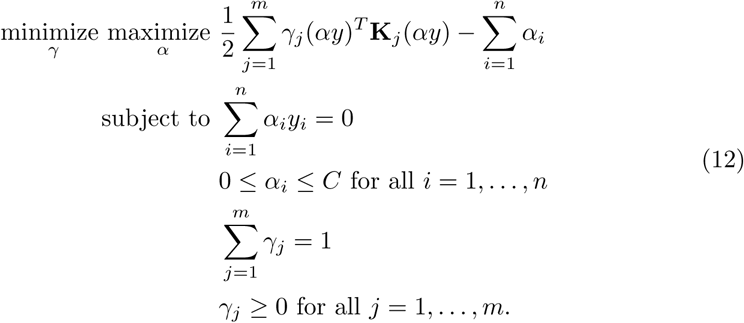

MKL allows is the flexibility to assign kernels on an individual variables basis, or as a data integration tool by assigning the different kernels to multiple data sources.

Differences in scale can be made even more dramatic with different kernel choices. An MKL algorithm can possibly put more importance on the variable with the largest variability regardless of accuracy of classification. Feature scaling is an important technique in many machine learning algorithms which transforms data so that they are on the same scale and unitless quantities. Common methods of data transformations are feature scaling, mean centering, or *z -* score transformation. Additionally, many articles have suggested dividing each kernel matrix by their respective trace to help speed up algorithms and eliminate computational issues. In some machine learning algorithms data normalization can reduce computation time, in SVM it can reduce the time to find support vectors and changes the classification rule.

There have been many algorithms proposed to conduct MKL. One class of MKL algorithms are wrapper methods which iteratively solve a single kernel learning problem for a given combination of kernel weights. Wrapper methods iteratively optimize *f, b, α* with *γ* fixed, sometimes referred to as the fixed weights problem, and then optimize *γ* with *f, b, α* fixed. A theme of wrapper methods is that they reformulate either the dual or primal of the MKL problem in order to use off-the-shelf efficient solvers. Bach *et. al* (2004), reformulated the quadratically quadratic programming problem (10) as a second-order cone programming (SOCP) problem. Bach uses sequential minimization on a smoothed version of QCQP, unfortunately SOCP with many samples can be quite slow. Sonnenburg *et. al* (2006) recast the MKL problem as a semi-infinite linear program (SILP) problem, which they propose column generation as a technique to solve the SILP. They also explore generic loss functions such as soft margin loss, one class margin loss, and *ϵ*–sensitive loss. Sonnenburg pointed out that a shortcoming of wrapper methods is optimization of *f, b, α* is inefficient, and unnecessary, if *γ* is not optimal and made recommendations to help MKL algorithms be efficient on large scale problems. Rakotomamonjy *et. al* (2008) develop a smooth formulation, with *L*2 regularization,of MKL that is equivalent to (9) by replacing 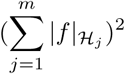 with 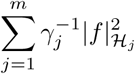 in the objective function. This relationship is established by utilizing the Cauchy Schwarz inequality to show the following relationship:

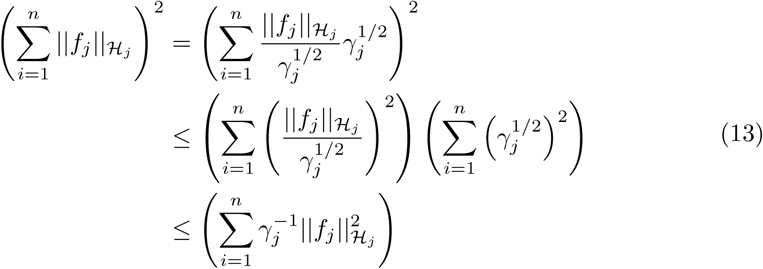

with equality when *f* and *γ* are linearly dependent, and in particular if

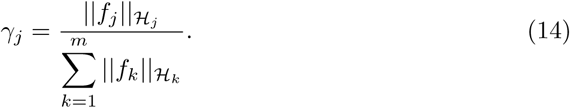

This is novel because it gives an explicit formula for makes the optimization less complex.

Bach et al. (2008) explored the Block 1-norm in order to compute regularization paths for multiple kernel learning. Bach points out the that Block 1-norm is a similar formulation to Lasso, but sparsity is enforced so that only a small number of kernels are used in the final model. Group lasso divides predictor variables into blocks and determines which blocks are most important for prediction, then assigns them the same regression coefficient. Group lasso can be formulated as:

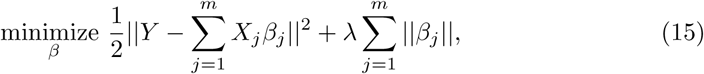

where *λ* ≥ 0, and *β*_*j*_ is the regression coefficient corresponding the *j*^*th*^ group of predictors. Group lasso reduces the number of parameters that need to be estimated in a model by requiring that each member of a group has the same regression coefficient, but there is no feature selection within groups. There have been further modifications to incorporate variable selection within groups, as well as, penalties that adjust for the size of a group.

Kloft *et. al*, and Xu *et. al*, both utilized the similarity of MKL and group lasso. Kloft noted that using the *L*1 rarely outperforms using uniform weights in MKL, and proposed an additional constraint in search of the best trade off between sparsity and uniform weights. Kloft showed the Tikhonov regularization:

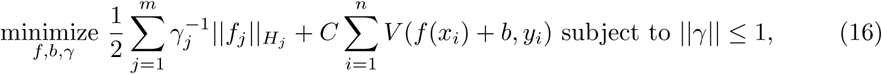

which is similar to Lasso regression formulation, and Ivanov regularization:

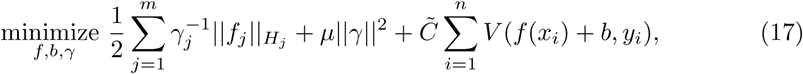

are equivalent. Incorporating the hinge loss into the Tikhonov regularization and taking dual lead to a similar quadratic programming problem to (10) which is solved with Newton’s method or a cutting plane algorithms. They propose an explicit formula,

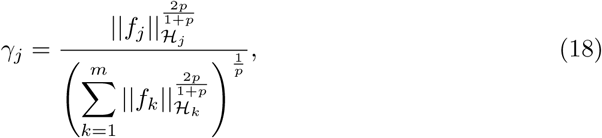

to compute optimal coefficients. Xu *et. al* exploits the relationship between group lasso and MKL. They provide an algorithm that is faster than Kloft’s algorithm and also does not rely on Taylor’s series expansion required for Newton’s method by applying (14) at each step as opposed to computing the weights with another optimization problem.

Suzuki *et. al* (2011) further explored block 1-norm proposed by Bach. The block 1-norm problem is solved using dual augmented-Lagrangian. This problem is solved by alternating between solving the primal problem using proximal descent and then solving the dual problem using Newton’s method. This novel method does not rely on iteratively solving linear or quadratic programming problems and can efficient solve MKL with more than 1000 candidate kernels. The paper provides a detailed overview of the derivation and outlines many scenarios that use elastic net and block *q* norm, for *q* ≥ 1, regularizations, as well as, logistic, squared, hinge, and *ϵ* – loss functions. These scenarios make DALMKL a promising method for extension to other employed in other arenas such as causal inference, or survival analysis.

SVM can easily be extended to support vector regression (SVR). The goal of SVR is to construct a prediction that is at most away from the observed response. The loss function is called *ϵ* – sensitive loss, and is defined as *ξ*_*i*_ = max(0, *‖ f* (*x*_*i*_) − *y*_*i*_ − ‖*ϵ*). Soft margin SVR is typically formulated as:

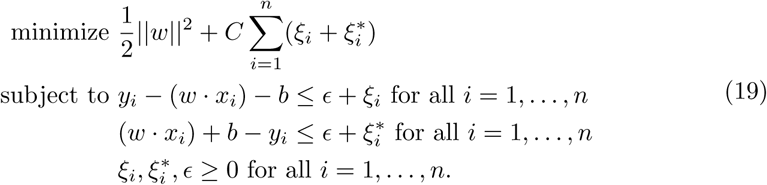

Here *ξ*_*i*_ and 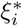 correspond to slack variables for the predicted values that are above or below the observed value respectively. This problem is not solved this problem directly, but rather we solve the dual form, given below:

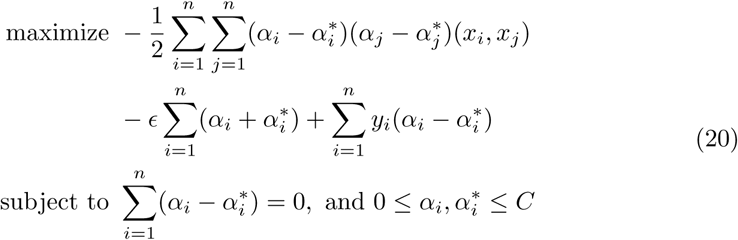

There are only subtle modifications, from derivations SVM and application of the kernel trick, required to solve this with quadratic programming solvers. We can use cross validation to find the optimal *C* and *ϵ*. Similar to before, larger *C* tend to smaller *ϵ*, large errors are heavily penalized. While small values *C* allow for larger *ϵ* since there is not a large penalty for large prediction errors. Fortunately, the tools developed in previous discussion can be utilized, namely we can extend SVR to non-linear regression by using the kernel trick. Taking the Lagrangian dual leads to quadratic programming problem, which can be solved using off the shelf solvers. A more detailed discussion of SVR can be found in Smola et al. (2004). As mentioned above, many authors have discussed that their MKL algorithms can easily be extended regression problem with.

## 3 RMKL R package

RMKL provides several methods to implement MKL, SVM, and multiple kernel regression. There are several dependencies of this packages. RcppArmadillo allow for “Seamless R and C++ Integration”, kernlab is used to create the kernel matrices, and caret is used to train SVM models.

Traditionally, kernel matrices are generated before implementation of MKL. A wrapper function is provided that can produce kernel matrices for both the test and training sets. Currently, RMKL can construct linear, polynomial, Gaussian, and clinical kernels proposed by Daemon.

Before implementing MKL, we recommend conducting SVM, with cross validation for two reasons. First, if SVM can successfully classify data, then there an opportunity for improving the accuracy using MKL. Secondly, if MKL is used as a data integration method, SVM can provide intuition for which parameters lead the best separation of data. To perform SVM, we have included a wrapper function that performs *k*-fold cross validation.

SimpleMKL has been implemented in previous cancer research studies. For example, Sloane *et al.* (2014) implemented SimpleMKL treating gene pathways as different data sources. SEMKL and DALMKL are more computationally efficient than SimpleMKL. SEMKL directly computes the kernel weights, while SimpleMKL uses gradient descent at each iteration to compute the kernel weights. DALMKL is written in C++, and uses Newton descent to update the kernel weights. Additionally, DALMKL has performs well when thousands of candidate kernels are used, and is dramatically faster than SimpleMKL. SimpleMKL and DALMKL tend to give sparse solutions allowing for an opportunity to interpret the final results.

We are not able to directly compare the performance of SimpleMKL and SEMKL using the same parameterization, however this is not true for DALMKL. Suzuki points out the relationship between the cost parameter of DALMKL and other methods, This conversion of cost parameterizations is available in the MKL package. Suzuki provided the following relationship

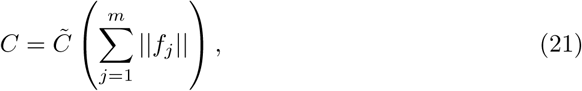

where *C* corresponds to the cost of misclassification of SEMKL or SimpleMKL, 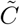 corresponds to cost of misclassification in DAMKL, and ‖ *f*_*j*_ ‖ = *α*^*T*^ **K**_*j*_*α*

Even though sparse MKL solutions do not typically outperform uniformly weighted kernels. there is still value in sparse kernel weights, specifically the model can be easier to interpret with fewer non-zero kernel weights. SimpleMKL tends to provide sparse solutions compared to SEMKL, SEMKL rarely produces kernel weights that are identically zero. This is due to the additional line search step for each update of the kernel weights. Xu provides details on implementing MKL using with different methods for updating the kernel weights. By ensuring that the vector of kernel weights was unit length under the *L*2 norm (which is more kin to ridge regression), as opposed to the kernel weights having unit length under the *L*1 norm (similar to lasso or group lasso variable selection schemes). DALMKL enjoys this flexibility too.

## 4 Results

### 4.1 Benchmark Example

Besides accuracy, an additional important characteristic of MKL is the selection of kernel weights. In this example, 9 datasets are generated, each dataset has two groups and the amount overlap between the two groups varies. These datasets can be generated RMKL. When there is no overlap, we expect that a radial kernel with a small sigma parameter to have a larger weight than a radial kernel with a larger sigma parameter (see kernel parameterization in the kernlab R package), leading to a smooth boundary. On the other hand, if there is a large amount of overlap between the groups, not only do we expect lower accuracy but also a classification rule that is more variable, thus a larger sigma hyperparameter should be preferred.

On each of the 9 datasets, two radial kernels, *K*_1_ and *K*_2_ with hyperparameters *σ* _1_ = 2 and *σ* _2_ = 0.04 respectively, were used. In figure 1a, notice that as amount of overlap between the two groups increases the weight for *K*_1_ increases providing a classification rule that is less smooth to accommodate for the overlapping groups. When there is little overlap between the groups, we see that *K*_2_ is given much more weight then *K*_1_, leading to a smooth classification rule for data that are perfectly separable. Note that SimpleMKL and SEMKL had nearly identical performance. Figure 1b displays the accuracy of the methods as the overlap between the groups changes. Notices that all algorithms can classify perfectly when there is no overlap, but when the groups are completely overlapping, the prediction accuracy of each algorithm is approximately 0.5.

**Figure 1.**
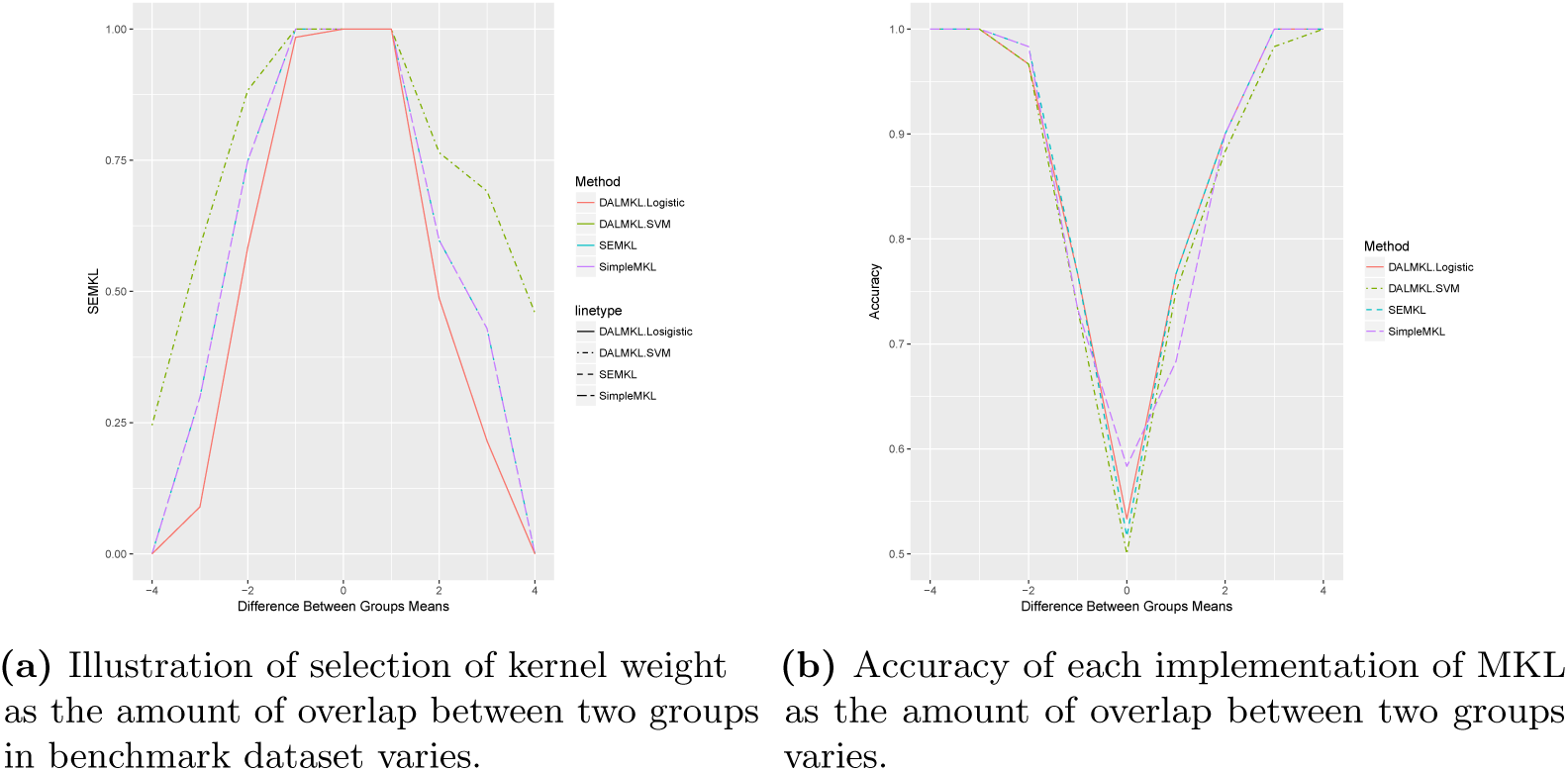
Results from SEMKL, SimpleMKL, and DAMKL on 9 benchmark datasets.

**Figure 2.**
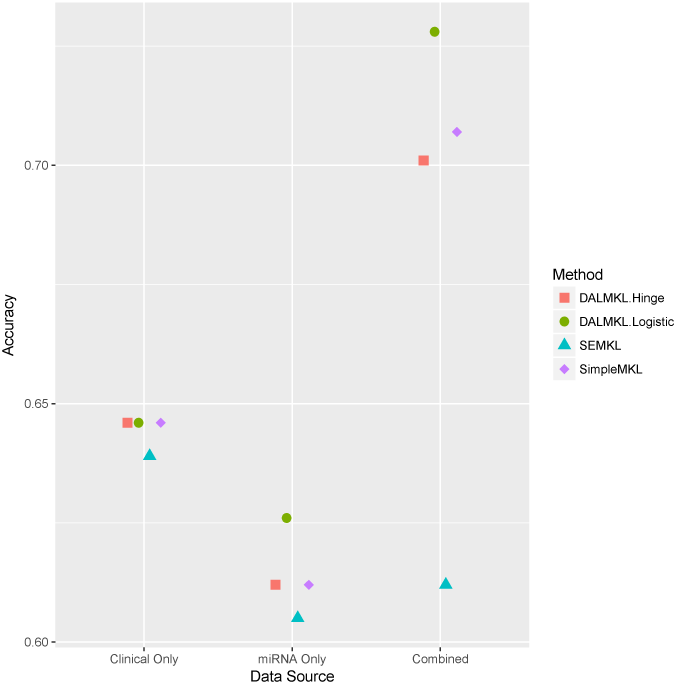
Comparison of performance of different MKL implementation using clinical data only, miRNA data only, and both clinical and miRNA data in a single analysis.

### 4.2 TCGA Ovarian

Survivorship for ovarian cancer is difficult to predict from clinical information only, which is limited since most cancers are late stage and occur only in females. Thus, information from high throughput data sources must be utilized to increase prediction accuracy. To illustrate MKL as a data integration tool, we use TCGA ovarian cancer data which contains both clinical and molecular profiles, and other data sources, from 572 tumor samples in total. Our goal is to use data from multiple platforms to predict if a patient will live longer than three years after diagnosis.

Clinical kernels were constructed using kernels for stage (nominal) and age (ordinal), and the average of these two as a kernel. Sloane *et al.* (2014) suggests using effect size or p-values to prioritize genes to include into kernels. We include the 65 top ranked genes, based on p-value from a t-test. Figure 1c illustrates the accuracy of the four algorithms using clinical data only, miRNA data only, and a combined analysis. Surprisingly, using miRNA data only leads to the worst performance, but using both data sources leads to a substantially higher accuracy than either of the individual data sources.

## 5 Conclusion

MKL is better suited for integration of multiple high throughout data sources than SVM, which is limited to a single kernel for the entire analysis, regardless of the number of features and number of data sources. Our simulations show that each of the three implantation of MKL perform similarly. However, using TCGA data, DAMKL with logistic loss tends to produce the most accurate prediction, which is consistent with Suzuki’s findings. SEMKL is consistently worse than the other methods. The poor performance of SEMKL may be due to kernel weights not being as sparse as other methods. TCGA data illustrates the opportunity to combine multiple data sources into a single analysis and substantially boost accuracy. RMKL is a useful tool for integrating high throughout data.

In the future, we hope to extend the RMKL package to several loss functions, specifically for applications in survival analysis setting.

